# A highly contiguous genome assembly of *Cyclamen persicum* to accelerate functional genomics and breeding

**DOI:** 10.64898/2026.07.17.739093

**Authors:** Kenta Shirasawa, Yusuke Akita, Yuki Mizunoe, Takejiro Takamura

## Abstract

*Cyclamen* is an economically important ornamental plant widely cultivated for its diverse floral characteristics and adaptation to cool climates. Despite its horticultural significance, genomic resources for this species remain limited, hindering molecular studies and genomics-assisted breeding. Here, we report the first highly contiguous nuclear genome assembly of *C. persicum* generated using high-fidelity long-read sequencing. The assembled genome spans 1.48 Gb, consisting of 126 contigs with an N50 length of 52.3 Mb. Telomeric repeat analysis identified eight contigs containing telomeric sequences at both ends, suggesting the presence of near-complete chromosome assemblies. Genome completeness assessment using BUSCO indicated 98.1% completeness. Repetitive sequences occupied 82.9% of the assembly, with long terminal repeat retrotransposons accounting for 42.1% of the genome. A total of 40,223 protein-coding genes were predicted, with a complete BUSCO score of 95.7%. Comparative orthogroup analysis with five representative eudicot species identified 430 orthogroups specific to *C. persicum* and 363 orthogroups shared exclusively between *C. persicum* and *Primula kwangtungensis*, indicating the presence of both lineage-specific and Primulaceae-conserved gene families. These findings provide critical insights into gene family evolution within Primulaceae and establish an essential comparative framework for future genomic studies. The genome resource presented here provides an invaluable foundation for investigating genome evolution, gene function, and trait-associated loci in cyclamen, effectively facilitating molecular breeding and genetic improvement in this ornamental species.

## 1. Introduction

Understanding plant genome structure, evolution, and functional diversity is a central goal of plant genomics. Over the past two decades, advances in high-throughput sequencing technologies and bioinformatic methodologies have enabled the generation of reference genomes for a wide range of angiosperms^1,2^, thereby accelerating comparative genomics and functional studies^3^. However, many horticulturally important ornamental plants remain underrepresented in genomic databases. This is often due to their complex genomic features, such as large genome sizes and high proportions of repetitive elements, thus limiting a molecular-level understanding of the traits that underpin their economic and biological value.

The genus *Cyclamen* (Primulaceae) comprises approximately 25 species distributed across Europe, North Africa, and the Middle East (https://powo.science.kew.org). Species of *Cyclamen* are widely cultivated as ornamental pot and garden plants and are characterized by diverse floral morphologies, vivid flower colors, tuberous growth habits, and adaptation to cool climates^4,5^. Although the representative diploid chromosome number in the genus is 2*n* = 48, *Cyclamen* species exhibit substantial variation in genome size, chromosome number, and ecological niche, making the genus an intriguing system for studying genome evolution and speciation within the Primulaceae^5–8^. However, the genomic mechanisms driving this structural and size variation, as well as the dynamics of repetitive elements, remain largely unresolved due to the lack of high-quality reference genomes. Among the genus members, *Cyclamen persicum* is of particular importance, as nearly all modern cultivated cyclamen varieties have been developed from this species. Cultivars of *C. persicum* display extensive variation in flower color and morphology, making the species a valuable system for studying the genetic basis of ornamental traits. Thus, the establishment of a high-quality reference genome for *C. persicum* will not only advance our understanding of genome evolution within *Cyclamen* but also provide a cornerstone for future pan-genomic and evolutionary studies across the genus.

Molecular biological studies in *Cyclamen* have increased in recent years. Several studies have identified genes involved in flower color, floral organ development, and related traits^9–18^. Additionally, molecular resources have been generated through chloroplast genome sequencing and RNA sequencing–based transcriptome analyses^19–22^. While these studies have provided useful insights, they have been conducted largely at the single-gene or transcriptome level, without a structural genomic context. Consequently, they offer limited information on genome organization, repetitive element dynamics, and the evolutionary context of gene families. Moreover, the lack of a reference nuclear genome has hindered comprehensive genomic analyses and the high-throughput development of molecular markers in *Cyclamen*. A highly contiguous genome assembly of *C. persicum* will provide a framework for systematic gene annotation, investigation of genome structure, and comparative analyses within the Primulaceae and related taxa.

Plant genome assembly, however, poses significant challenges, particularly in perennial ornamental species characterized by large genome sizes, high heterozygosity, and complex genome architectures shaped by repetitive elements and structural variation. In recent years, whole-genome sequencing has been increasingly applied to horticultural plants with such complex genomes, driven by rapid advances in long-read sequencing technologies with high base accuracy, such as PacBio HiFi^23,24^. These advances now enable the generation of near-complete, highly contiguous assemblies even for species with intricate genomic structures^25,26^. Furthermore, the emergence of deep-learning-based gene prediction workflows has facilitated highly accurate, *ab initio* gene prediction even in the absence of comprehensive transcriptomic resources^27^.

In this study, we report a highly contiguous reference genome assembly of *C. persicum* generated using PacBio HiFi long-read sequencing. This high-quality genome assembly represents the first comprehensive nuclear genomic resource for the genus *Cyclamen,* providing a foundational platform to address evolutionary questions within the Ericales and establishing a robust molecular resource to accelerate breeding and functional genomics in this highly valued ornamental crop.

## 2. Materials and methods

### 2.1. Plant materials

Young leaves of *C. persicum* ‘KN purple’, cultivated in a greenhouse at Faculty of Agriculture, Kagawa University, were collected for genome DNA extraction. The ‘KN purple’ plants had been propagated by repeated inbreeding including self-pollination.

### 2.2. Genome size estimation

To construct a short-read library, a PCR-free library was prepared using the Swift 2S Turbo Flexible DNA Library Kit (Swift Biosciences, Ann Arbor, MI, USA). This library was then converted into a DNA nanoball sequencing library using the MGI Easy Universal Library Conversion Kit (MGI Tech, Shenzhen, China) and sequenced on a DNBSEQ G400 instrument (MGI Tech) to generate 2 × 100-bp paired-end reads. The genome size of *C. persicum* was estimated by *k*-mer frequency analysis (*k* = 17) using Jellyfish^28^ v2.3.0 (*k*-mer size = 17).

### 2.3. Genome assembly

Genomic DNA was extracted from young leaves using the Genomic Tip (Qiagen, Hilden, Germany). The genomic DNA was sheared to an average fragment size of 30 kb using Megaruptor 2 (Diagenode, Liege, Belgium) in the Large Fragment Hydropore mode. The sheared DNA was then used for HiFi SMRTbell library preparation using the SMRTbell Express Template Prep Kit 3.0 (PacBio, Menlo Park, CA, USA). The resultant library was separated on BluePippin (Sage Science, Beverly, CA, USA) to remove short DNA fragments (<15 kb) and sequenced using SMRT Cell 8 M on the Sequel II system (PacBio).

The obtained HiFi reads were assembled using Hifiasm^29^ v0.16.1 with default parameters. Organelle genome sequences, identified by sequence similarity searches of the reported plastid and mitochondrial genome sequences of cyclamen^2^ and primula^30^, respectively, using Minimap2^31^ v2.24, were eliminated. Assembly completeness was assessed with embryophyta_odb10 lineage data using Benchmarking Universal Single-Copy Orthologs (BUSCO)^32^ v5.2.2. Telomere sequences containing repeats of a 7-bp motif (5□-TTTAGGG-3□) were searched using the search subcommand of Tidk^33^ v0.2.1 with a window size of 1000 bp.

### 2.4. Gene and repeat prediction

Protein-coding genes were predicted using the *ab initio* strategy of Helixer^27^. The completeness of the predicted gene models was evaluated with the embryophyta_odb10 data using BUSCO^32^ v5.2.2. Repetitive sequences in the assembly were identified using RepeatMasker v4.1.6 (https://www.repeatmasker.org), based on repeat sequences registered in Repbase and a *de novo* repeat library built with RepeatModeler v2.0.4 (https://www.repeatmasker.org).

### 2.5. Comparative genomics analysis

Comparative orthogroup analysis was conducted using OrthoFinder v2.5^34^ with predicted protein sequences from *C. persicum* and five representative eudicot species, including *Arabidopsis thaliana* (Araport11, https://www.arabidopsis.org)^35^, *Vitis vinifera* (12X, https://www.genoscope.cns.fr/vitis)^36^, *Camellia sinensis* (CsSME v1.0, GenBank accession number of GCA_041154535.1)^37^, *Primula kwangtungensis* (Prkwt_v. 1.0, GCA_054094925.1)^38^, and *Rhododendron ripense* (RRI_r1.1, GCA_019656295.1)^3^. Orthologous gene groups shared among species and species-specific orthogroups were visualized using an UpSet plot^39^.

## 3. Results

### 3.1. Genome assembly and annotation

To estimate the *C. persicum* genome size, clean short-read data (70.4 Gb) were subjected to *k*-mer distribution analysis. The results indicated that the *C. persicum* ‘KN purple’ genome exhibited low heterozygosity, with an estimated haploid genome size of 1.8 Gb (Figure 1). A total of 2.9 million HiFi reads (41.3 Gb; 22.9× coverage of the estimated genome size), were obtained from three SMRT cells and assembled into 798 contigs. A total of 672 potential contaminant sequences (32.4 Mb in total) from organelle genomes were removed. Consequently, 126 contigs totaling 1.48 Gb, with an N50 of 52.3 Mb, were obtained (Table 1). These 126 contigs represented the genome assembly of *C. persicum* and were designated CPE_r1.0. Notably, more than 95% of the assembled sequence (1.41 Gb) was represented by 40 contigs (Figure 2). Telomere repeats were found at both ends of 8 contigs and at one end of 26 contigs (Figure 2), suggesting that these 8 contigs were assembled at the complete telomere-to-telomere chromosome level. A total of 40,223 protein-coding genes were predicted from the genomic sequence (Table 1). Repetitive sequences occupied 1.23 Gb (82.9%) of the genome, of which LTR retroelements (624.1 Mb; 42.1%) represented the dominant repeat type (Table 2). The accuracy of the genome assembly and gene predictions were supported by BUSCO scores of 98.1% and 95.7%, respectively (Table 3).

**Figure 1.**
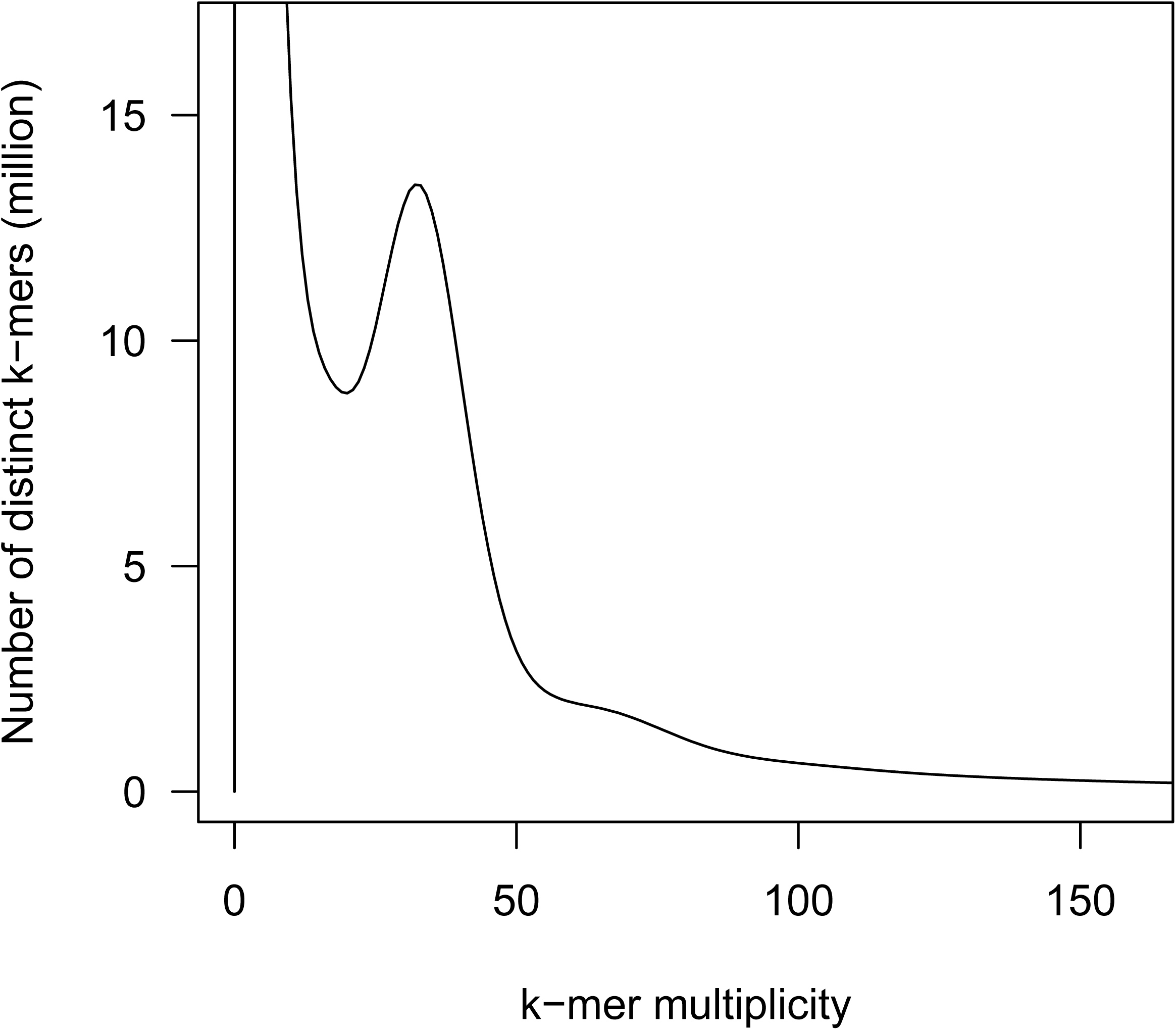
Estimated genome size of the *Cyclamen persicum*, based on *k*-mer analysis (*k* = 17), with the given multiplicity values.

**Figure 2.**
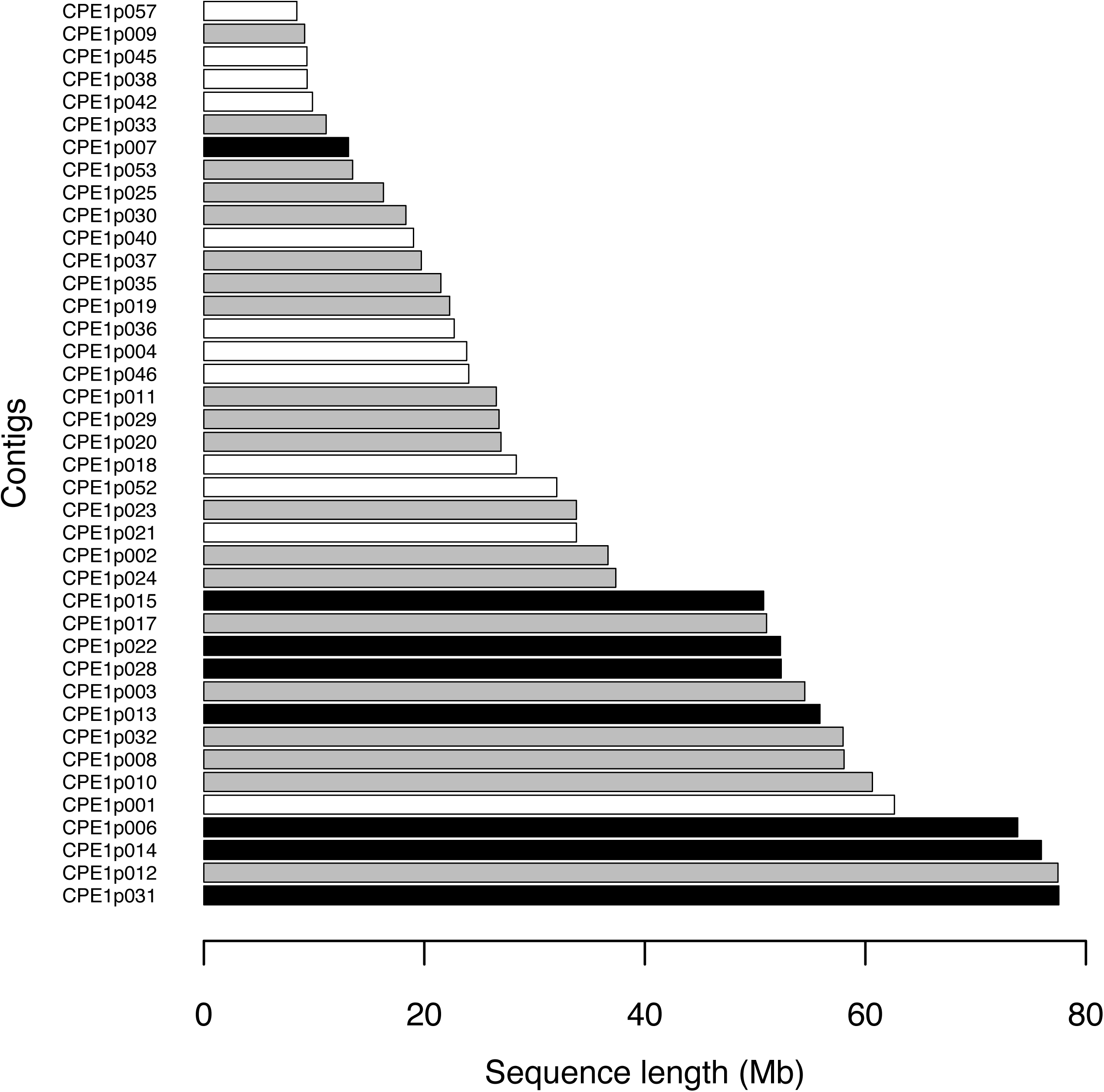
Contig length distribution for the top 95% of the genome assembly. Black, gray, and white bars represent contigs with telomeric motifs at both ends, at one end, or with no detected motifs, respectively.

**Table 1.**
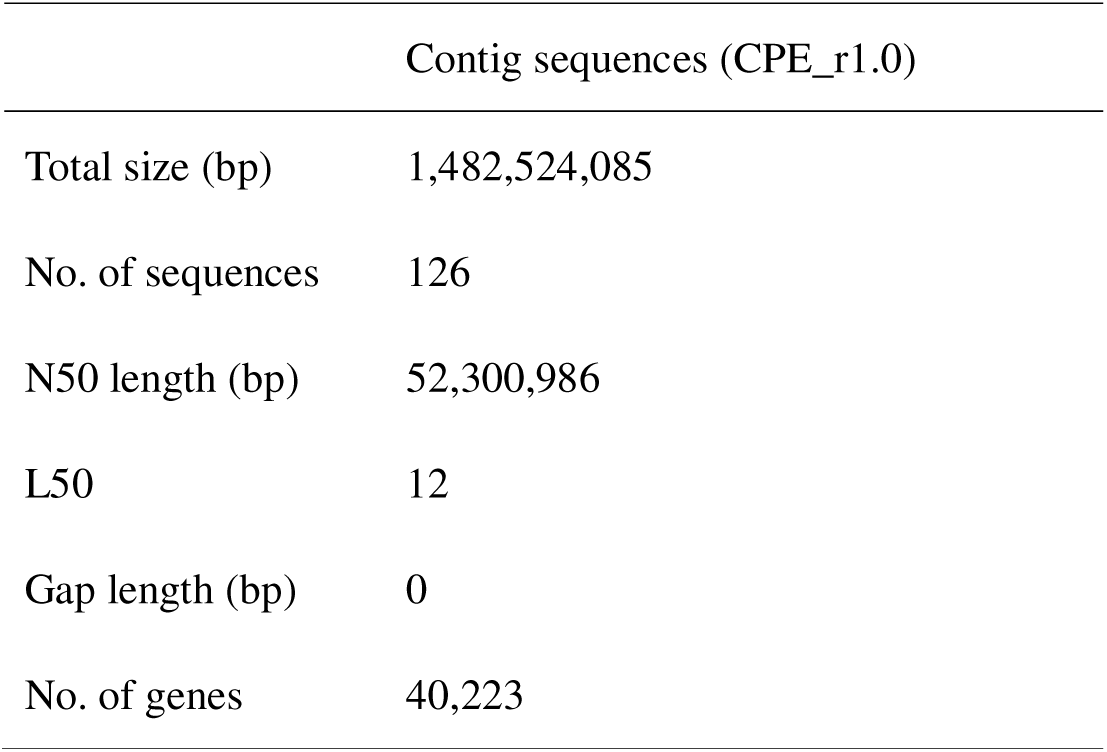
Assembly statistics of the *Cyclamen persicum* genome.

**Table 2.** Repetitive sequences in the cyclamen genome.

| Repeat type | No. of repeats | Total length (bp) | Relative proportion (%) |
| --- | --- | --- | --- |
| SINEs | 5,876 | 535,801 | >0.0 |
| LINEs | 32,393 | 18,568,281 | 1.3 |
| LTR elements | 395,762 | 624,119,859 | 42.1 |
| DNA transposons | 85,726 | 40,404,166 | 2.7 |
| Small RNA | 2,776 | 4,042,171 | 0.3 |
| Satellites | 8,922 | 4,314,249 | 0.3 |
| Simple repeats | 226,076 | 13,643,877 | 0.9 |
| Low complexity | 25,922 | 1,267,119 | 0.1 |
| Unclassified | 1,154,127 | 520,501,486 | 35.1 |

**Table 3.** Completeness of the genome assembly and predicted genes.

| BUSCOs | Genome assembly (%) | Predicted genes (%) |
| --- | --- | --- |
| Complete | 98.1 | 95.7 |
| Single-copy complete | 90.2 | 88.2 |
| Duplicated complete | 7.9 | 7.5 |
| Fragmented | 0.7 | 2.4 |
| Missing | 1.2 | 1.9 |

### 3.2. Comparative orthogroup analysis

To investigate gene family conservation and diversification in *C. persicum,* comparative orthogroup analysis was conducted using the predicted protein sequences from *C. persicum* and five representative eudicot species: *Arabidopsis thaliana, Vitis vinifera*, *Camellia sinensis*, *Primula kwangtungensis*, and *Rhododendron ripense* (Figure 3).

**Figure 3.**
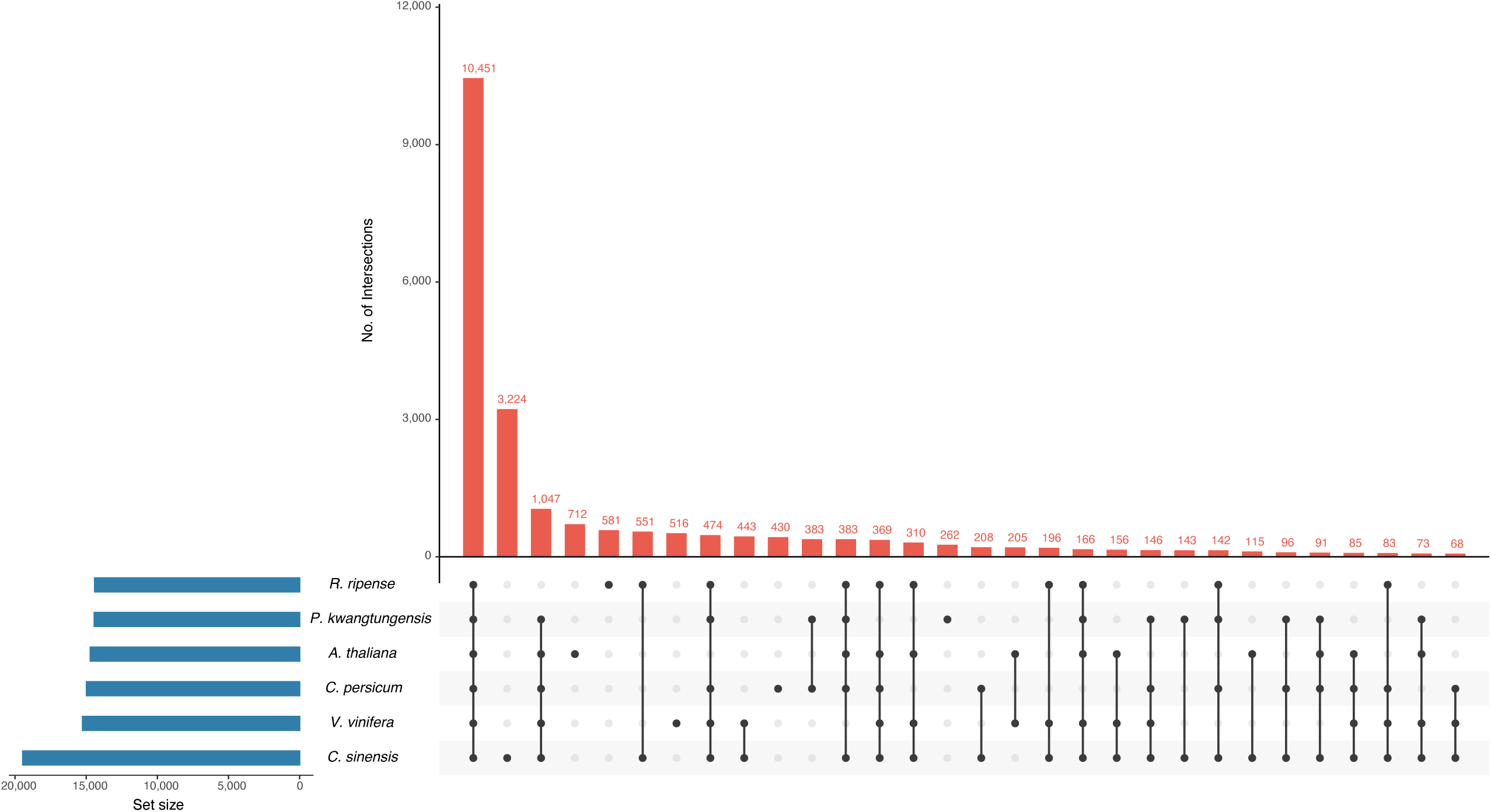
Comparative orthogroup analysis among *Cyclamen persicum* and five representative eudicot species. Shared and species-specific orthogroups were identified using OrthoFinder and visualized as an UpSet plot. Numbers indicate orthogroups shared among specific combinations of species.

Orthogroup analysis revealed extensive conservation of gene families among the examined species, with numerous orthogroups shared across multiple eudicot lineages. Notably, 430 orthogroups were identified exclusively in *C. persicum*, indicating the presence of lineage-specific gene families that may have arisen during the evolution of *Cyclamen*. In addition, 363 orthogroups were shared exclusively between *C. persicum* and *P. kwangtungensis*, the two representatives of Primulaceae included in this study, suggesting the retention of gene families that are characteristic of the family. The analysis also identified 474 orthogroups shared among all examined species except *A. thaliana*, and 1,047 orthogroups shared among all species except *R. ripense*. These patterns may reflect lineage-specific gene loss, divergence, or differences in genome annotation among species. Overall, the results indicate that the *C. persicum* genome contains both highly conserved eudicot gene families and lineage-specific gene repertoires, providing a useful framework for future comparative and evolutionary studies within Primulaceae and Ericales.

## 4. Discussion

In this study, we generated a highly contiguous reference genome assembly of *C. persicum* (CPE_r1.0), resolving the 1.48 Gb genome into just 126 contigs with a contig N50 length of 52.3 Mb (Table 1). The discrepancy between the estimated genome size (Figure 1) and assembly length is likely attributable to collapsed highly repetitive regions (Table 2) and/or overestimation in *k*-mer analysis^40^. The high BUSCO completeness score (98.1%) confirms that most of the conserved gene space was successfully captured (Table 3). Although *C. persicum* generally exhibits strong inbreeding depression^41^ and high heterozygosity in cultivated lines, our *k*-mer analysis revealed relatively low heterozygosity in the targeted accession (Figure 1). This indicates that the strategic selection of a partially inbred line successfully resolved the computational haplotype-purging challenges typical of outcrossing species, demonstrating the effectiveness of high-accuracy PacBio HiFi long-read sequencing for complex plant genomes.

Repetitive sequences occupied 82.9% of the CPE_r1.0 assembly, dominated by LTR retrotransposons (Table 2). Notably, 35.1% of these repeats remained unclassified, suggesting a substantial reservoir of lineage-specific elements that likely drove genomic expansion in *Cyclamen*, consistent with other large-genome angiosperms^42^. Despite this heavy repeat burden, our annotation using the *ab initio* pipeline of Helixer successfully identified 40,223 protein-coding genes (Table 1) with a high BUSCO completeness of 95.7% (Table 3). This level of completeness, achieved without transcriptome-assisted training, confirms that CPE_r1.0 provides robust and reliable gene models suitable for downstream comparative and functional analyses, with gene numbers aligning well with those of related eudicots^43,44^.

Our comparative orthogroup analysis highlighted 363 gene families shared exclusively between *C. persicum* and *P. kwangtungensis*, representing core family-specific synapomorphies conserved within the Primulaceae family (Figure 3). These gene families provide an important starting point for identifying traits conserved within Primulaceae. Furthermore, the 430 *C. persicum*-specific orthogroups represent prime candidates underlying genus-specific traits, such as unique floral ontogeny, tuberous growth habits, and specialized volatile profiles. CPE_r1.0 now serves as a critical genomic resource to integrate fragmented transcriptomic data^19–22^ and leverage established transformation systems^45,46^ to facilitate transgenic phenotypic modifications^9,47^. From an applied perspective, CPE_r1.0 offers a valuable framework to overcome conventional breeding bottlenecks in this outcrossing species^48^, thereby accelerating genomics-assisted breeding and functional genomics in this ornamental crop.

## Acknowledgments

We thank Y. Kishida, C. Minami, K. Ozawa, H. Tsuruoka, and A. Watanabe (Kazusa DNA Research Institute) for technical assistance.

## Data availability

Raw sequence reads and assembled sequences are available at DDBJ (BioProject accession number PRJDB42699). The genome assembly and gene annotation files are available in Kazusa Genome Atlas (https://genome.kazusa.or.jp).

